# Spring growth variation in pioneer and fibrous roots in *Abies sachalinensis* seedlings from provenances with contrasting snow cover environments

**DOI:** 10.1101/2022.09.03.506439

**Authors:** Tetsuto Sugai, Wataru Ishizuka, Taiga Masumoto, Izuki Endo, Jun’ichiro Ide, Saki Fujita, Makoto Kobayashi, Naoki Makita

## Abstract

**Premise of research:** In boreal forests with various snow cover conditions, local adaptation can be expected in spring root exploration due to soil freezing or competition for the fundamental niche. However, few studies have investigated the adaptive strategies for acquiring soil space and resources after snow thawing. To get insights for the eco-evolutional significance of root growth dynamics in boreal regions, this study evaluated the intraspecific variation of morphology and structures in pioneer roots and fibrous roots.

**Methodology:** Validation model specie was *Abies sachalinensis*, a representative boreal conifer species with local adaptation. Five-year-old seedlings derived from an eastern provenance with low snow cover and a northern provenance with high snow cover were grown for two years in a common garden of high snow cover region. To predict spring growth patterns of roots, seedlings were dug up in early spring just after snow thawing and late spring after one month has passed. Pioneer roots exploring soil space and fibrous roots acquiring soil resource were respectively collected, and their functions were evaluated by morphology and structures indicating its construction costs and maturity.

**Pivotal results:** In pioneer roots, morphological and structural difference between sampling months were significant only in northern provenance, and high construction cost and well-developed structure were observed in early spring. This represents the intraspecific variation in reaction norms of pioneer roots to spring soil environment, indicating the soil space exploration in northern provenance may start in early spring or earlier. In fibrous roots, differences of specific root length and area between sampling months were significant only in eastern provenance, and low construction cost were observed in early spring. This may reflect the plastic responses of fibrous roots to the condition of high snow cover region.

**Conclusions:** This study indicated that local adaptation to contrasting snow cover condition may lead to the differentiation of spring root growth dynamics for the competitive or conservative strategies. Our results address better understandings of the mechanisms of niche acquisition in evergreen conifer.

## Introduction

In the cool-temperate zone, temporal variability of environments is recognized as a season. To the seasonal dynamic, a developmental pattern of traits is often adaptive (Ghalambor et al. 2007, Forrest and Miller-Rushing 2010). The phenological optima are expected to be specific to the regional environments, causing the intraspecific variation of phenology because of local adaptation (De Lisle et al. 2022). Since the intraspecific variation in phenology is the consequence of reaction norms to seasonal stimuli (West-Eberhard 2003), its patterns provide cues for predicting how organisms seasonally react to climate changes (Franks et al. 2014). In trees, the intraspecific variation in the development of above-ground traits has been elucidated. The significant intraspecific variation has been often observed in specific seasons (West-Eberhard 2003), such as the timing of bud burst (Mediavilla & Escudero 2009), the leaf longevity (De Kort et al. 2016), freezing tolerances (Noordermeer et al. 2021), but also in the seasonal pattern itself, such as photosynthetic activities (Fréchette et al. 2020, Sugai et al. 2023) and stem growth (Sanmiguel-Vallelado et al. 2021). However, the body of studies on the intraspecific variation in the below-ground dynamics is still growing yet, for example, the non-structural carbon (Wang et al. 2018), the growth pattern of fine roots (Ma et al. 2022). Moreover, little information is available regarding the root phenology (McCormack et al. 2015, Tamura et al. 2022, Liu et al. 2022). The insights derived from root phenology and its intraspecific variation would shed light on the ecological significance for understanding the mechanisms of niche acquisition and their evolutionary aspects (Begon et al. 2006).

Soil temperature has been considered as the driver of root development given sufficient precipitation (Ding et al. 2020). The sensitivity of fine root elongation to soil temperature was variable between tree species (Pregitzer et al. 2000). In evergreen conifer species, the relatively high quantity of fine roots has been recognized in spring (Persson 1978), when the fine root elongation rate of conifers was correlated only below 8 °C of soil temperature (Wang et al. 2018). This fine root elongation at relatively low soil temperatures, where other species including deciduous trees would not develop (Steele et al. 1997), may reflect the ecological characteristics as late successional species who can acquire niche early and stay in for the long term (Makoto et al. 2020). These studies indicated that the root phenology would have the potential to be evolved, and particularly boreal evergreen conifers would have developed the root dynamics with relatively low soil temperature dependence at the beginning of the growing season when low soil temperatures begin to increase in spring (Ding et al. 2020).

In soils with heterogeneous nutrient and physical properties, roots often prioritize exploration rather than acquisition (de Kroon & Visser 2003, Begon et al. 2006). Among the various root taxonomies, pioneer roots are visually and functionally distinguishable from fibrous roots (Figure 1). Compared with fibrous roots, pioneer roots have been known to have a thick, well-developed structure and requires high construction costs (Zadworny & Eissenstat 2011, Montagnoli et al. 2021) which lead to the tolerances to anaerobic conditions with superior transportability and the exploration function of soil space. On the other hand, fibrous roots often show a relatively thin diameter and cortex layers without lignified woody cells, which enable them to efficiently absorb soil water and nutrients, and symbioses with mycorrhizal fungi (de Kroon & Visser 2003). The clear sequential stages of development are recognized in these roots (Wilcox 1954), where pioneer roots develop first, and then fibrous roots often emerge laterally from the maturity zone of pioneer roots. However, the intraspecific variations in the growth patterns of these root types have not been verified simultaneously so far.

**Figure 1:**
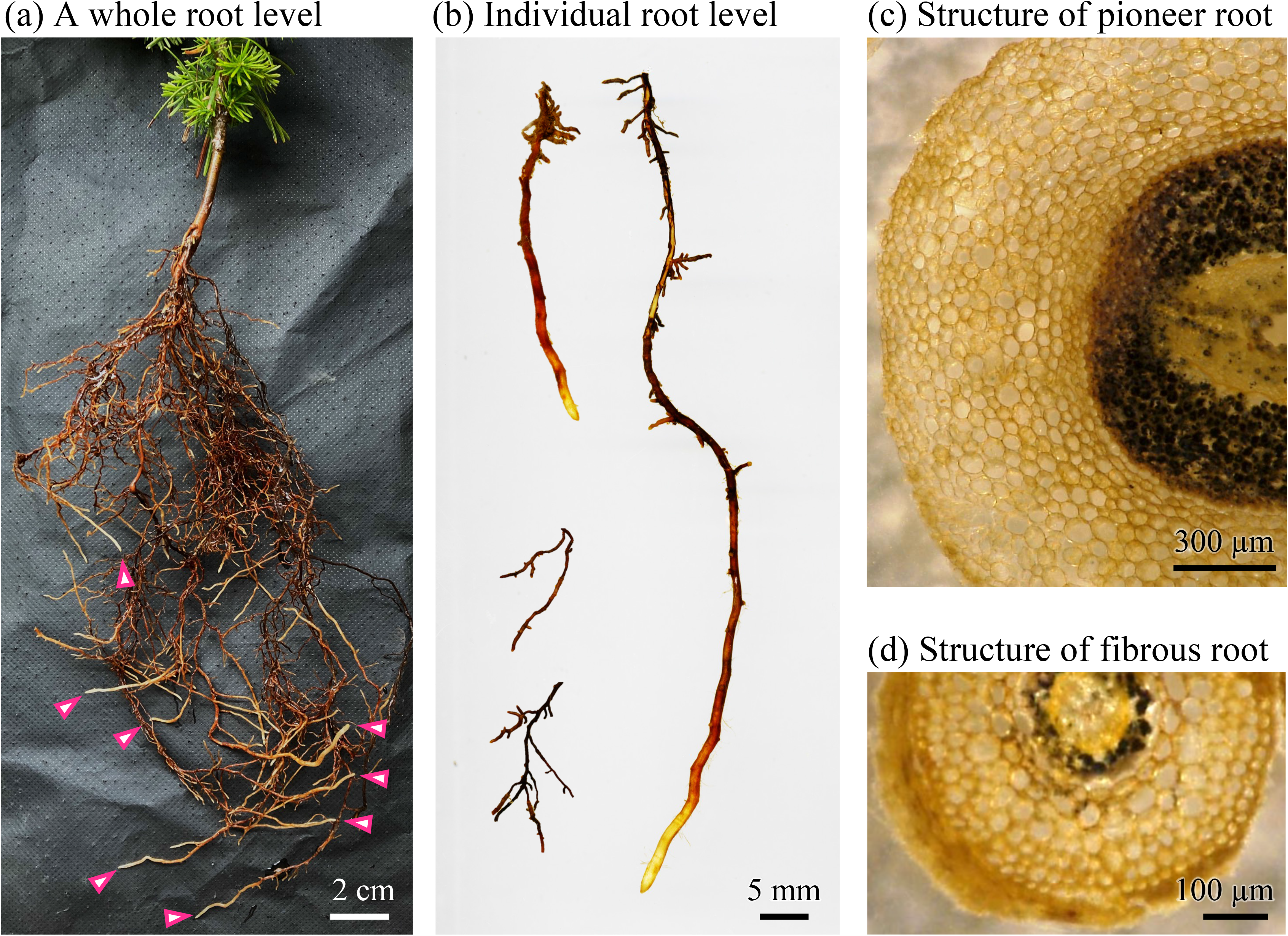
Diagram of representative pioneer roots and fibrous roots in seedlings of *Abies sachalinensis*. (a) Underground part of a seedling with pioneer roots denoted by pink arrowheads. (b) Individual pioneer roots with thick and white root tips and fibrous roots with thin and dark-brown root tips. (c-d) Section images showing internal structure of a pioneer root (c), and a fibrous root (d), which were prepared as a thickness of 50 µm and stained with iodine solution.

This study investigated the morphology and structure in different root types in a boreal evergreen conifer species, Sakhalin fir (*Abies sachalinensis*). In Sakhalin fir, the local adaptation has been reported mainly in above-ground traits (e.g., Ishizuka et al. 2021, Sugai et al. 2023), which would represent responsible association with winter climate environments. In boreal forests, the regional variation in spring soil temperature depends on snow cover conditions (Fukuzawa et al. 2021, Makoto et al. 2022). In high snow cover regions, soil temperature and resource availability would increase rapidly (Bardgett et al. 2005) more than low snow cover regions because soil suddenly faced relatively high air temperatures when the thermal interception by snow cover ends. In low snow cover regions, however, the rapid root elongation can lead to a risk of encountering soil freezing. We expected the fast development of pioneer roots would be adaptive for high snow cover provenances since the competition for underground resources in spring will be intense. If so, a relatively thick and well-developed pioneer roots may be observed in high snow cover provenance. On the other hand, fibrous roots branching later than pioneer roots may be dominated by seasonal stimuli without genetic variation to optimize the balance between construction costs and its benefit (de Kroon & Visser 2003). Here, we elucidated the intraspecific variation in the functional morphological traits; specific root length (ratio of root length to mass), specific root area (ratio of root surface area to mass) and root tissue density (ratio of root mass to volume), and the anatomical traits; central cylinder, cortex, and epidermis, which could exhibit its functions and the resource availability (McCormack et al. 2015, Yahara et al. 2019). We conducted a common garden trial, where seven-year-old seedlings originating from high and low snow cover provenances were dug up in early spring just after snow thawing and late spring after one month has passed to predict spring growth patterns of roots.

## Materials and methods

### Study site and material

A common garden trial was conducted at the nursery of the Forestry Research Institute, Hokkaido Research Organization, located at central Hokkaido (43°29N, 141°85ʹE; 45 m a.s.l.). As an experimental material, Sakhalin fir (*Abies sachalinensis*) was adopted in this study. Sakhalin fir distribution is on a northern island in Japan, Hokkaido in which various environmental conditions are recognized along with a geographic gradient, e.g., maximum snow depth from 20 cm to more than 400 cm mainly from eastern side to western side (Supplemental material Appendix 1). In the most regions of Hokkaido, snow begins to thaw in mid-March, and it has widely disappeared by early May. Previous studies have reported the genetic variation of Sakhalin fir in above-ground traits such as home advantage in growth performance, frost tolerance of winter buds and disease resistance (Sakai 1983, Eiga 1984, Ishizuka et al. 2021). In addition, boreal evergreen conifers, including Sakhalin fir, have a relatively early onset and late cessation of roots development from mid-April when covered snow just started melting to mid-December when snow have started covering soil (Satoh 1995). Thus, Sakhalin fir is a valuable model for testing the adaptive developmental patterns of different root types. In this common garden trial, open-pollinated progeny of selected mothers (family lines) were planted. Plant materials were sown in 2015 using seeds produced in a seed orchard of Sakhalin fir in 2014 and transplanted twice at spring of 2017 and 2019. Tested progeny were selected from 8 family lines derived from two provenances with contrasting environmental condition, i.e., (i) the eastern-edge provenance (43°07ʹ ± 0.01N, 145°03ʹ ± 0.01E) and (ii) the northern provenance (44°35ʹ ± 0.08N, 142°32ʹ ± 0.01E). At the time of second transplanting in 2019, the experimental plots of approximately 2.2 m × 2.7 m were set up with three replications in the same field with the growing field before transplanting, with seedlings planted approximately 50 cm apart in each. Basic meteorological information on the selected family lines and the cultivated experimental environmental conditions were summarized (Supplemental material Appendix 1, Supplemental material Appendix 2). After transplantation, seedlings were cultivated with commercial fertilizer and automatic irrigation during two growing seasons except from November to May during 2019–2020.

In a previous study, pioneer roots of Sakhalin fir seedlings were observed in late March under snow cover at the same site of this experiment (Satoh 1995), which was validated again in this study (Supplemental material Appendix 3). Based on them, seedlings were dug up both in April immediately after snow thawing, and in May when the soil temperature has increased sufficiently after one month has passed since the snow was thawing. A total of 48 seven-year old seedlings of seven-year-old (24 seedlings per month) were selected as samples for the following analysis. Then, 12 seedlings in northern provenance and 12 seedlings of eastern provenance were collected from 3 plots (4 seedlings of northern and eastern provenances per plot) in each month. Of these, three individuals were excluded from samples in May due to abnormal growth conditions, so the final number of seedlings collected was 45 in this study Additionally, the seedlings of both northern and eastern provenance continued to grow without being sampled in spring were collected in December 2021 and February 2022 (Supplemental material Appendix 3).

### Root morphology

Immediately after cutting the above-ground parts of each individual seedling, a below-ground part was carefully collected by shovels for each individual. The attached soil with a below-ground part was removed by tap water within the bucket which was kept for the following drymass measurement. After cleaning the whole root system, the length and number of pioneer roots in the below-ground part were measured. In this study, pioneer root was distinguished from other fibrous roots based on the diameter and white color from the elongation zone to maturity zone of a root (Figure 1, Supplemental material Appendix 4). From each individual, five pioneer roots were randomly selected from all the pioneer roots to measure the length of its root length, which would indicate the function to explore soil space and/or its maturity (Zadworny & Eissenstat 2011, Montagnoli et al. 2021). The length of the pioneer root was measured as the length from the root end to the elongation zone with bright white by a ruler to the nearest 1 mm. The number of all pioneer roots per individual was counted, including the number of this root in stored soil water in the bucket. After pioneer root measurements were completed, intact fibrous roots and pioneer roots were randomly collected from whole roots of individuals to evaluate morphological and anatomical characteristics, respectively. Finally, the roots were dried and used for drymass measurements, where all roots contained the stored soil were also used. The fibrous and pioneer roots collected for morphometrics were scanned with a flatbed scanner (GT-X980, EPSON, Tokyo, Japan) immediately after collection. The scanned images were analyzed using root image analysis software (WinRHIZO, REGENT Inc, Quebec, Canada) to obtain root length, surface area, volume, and average diameter. The dry weight of roots completed scans was measured by 0.1 mg increments. Then, specific root length (SRL, m g^-1^), and specific root area (SRA, cm^2^ g^-1^) were obtained as the ratio of dry weight to length and area of roots, and root tissue density (RTD, g cm^-3^) was obtained as the ratio of volume to dry weight of roots.

### Root anatomy

The fibrous and pioneer roots for evaluating tissue structure were impregnated with 2% glutaraldehyde solution immediately after collection. After one day of permeation of the fixative solution into the tissue, the tissue was stored in 80% ethanol solution. To prepare sections from the samples, the samples were initially traversed mainly from the maturation zone with a cutter, and a few centimeters of the sample were fixed horizontally on a freezing stage (MC-802A, YAMATO, Saitama, Japan). Sections with a thickness of 50 µm were cut on a microtome (REM-700, YAMATO, Saitama, Japan) and stained with iodine solution for approximately 30 seconds. The sections were then observed under an optical microscope and the tissue structure was captured (VH-5500, KEYENCE, Osaka, Japan). This process was repeated approximately three times per sample, and the image with the clearest structural characteristics among the images obtained was used for subsequent image analysis.

The image analysis of root sections was performed by a free image processing software (ImageJ, NIH, USA) in this study. As root tissue structures, diameter, cortex, and central cylinder were measured in the obtained section image process (Supplemental material Appendix 4). These characteristics were measured at three locations from a single image and their average value was calculated. Since the linear relationships between a diameter of root section and average size of each cell as well as the other anatomical values were observed, these anatomical values in each sample were converted to relative ones to the diameter of root section, respectively.

### Mass and structural fraction

For the pioneer root quantity at a whole plant level, total weight was estimated from SRL and total root length estimated from the total number and average length per seedling. All the sampled roots were dried in oven at 75 °C for approximately seven days and weighed. The roots used for the weight measurement were divided into woody roots with a diameter of 1.5 mm or more since the average diameter of the identified woody roots ranged up to approximately 1.5 mm in this study (Supplemental material Appendix 5). The other part except for woody root, where root branching was difficult to identify, was classified as a rootstock. The dry mass of fibrous roots was obtained by subtracting the estimated dry weight of the pioneer roots from the weights of the other roots except for woody roots and rootstock.

### Statistical analysis

All statistical analyses were performed by an open-source package for statistical computing (R ver.4.1.2, R Core Team 2019). In this study, missing values accounted for maximum of approximately 30% in the data of pioneer roots and a minimum of approximately 8% in fibrous roots. Since the complete case analyses with the process of excluding missing values should led to biased (Ellington et al. 2014), we alternatively adopted the multiple imputation method using the *mice* package (Nissen et al. 2019). The effects of sampling months and the provenances (i.e., genetic variation) on the following traits; total root dry mass, mass fraction of each root types, average length and total number of pioneer roots, SRL, SRA, RTD, average diameter, cortex, and central cylinder were tested by Two-way ANOVA in the following generalized linear mixed model. Because of the unbalanced data structure in this study, a type III variance analysis was performed using the *car* package. For this analysis, all values were used after logarithm transformation. The modeling was conducted by *lme4* package. The model was constructed as follows; where *Y_jkl_* denotes the measured values of traits of progeny of *k*-th family line derived from *j*-th region at *i*-th sampling months and located in *l*-th plots. *M_i_* denotes the effect of sampling months, *G_i_* denotes the effect of provenances, and *M_i_ × G_j_* denotes the interaction effect of sampling months and provenances. For the random effects, *ε_ij_, ε_k_*, and *ε_l_* denote the family lines of progeny, the plot replications, and error term, respectively. To evaluate differences between any of the groups, the multiple comparison adjusted by Tukey method was conducted as a post-hoc test using the *emmeans* package. A significant level of 5% was adopted for all statistical tests in this study.

## Results

While any significant changes were not observed in total root dry mass, the mass fraction of pioneer roots were significantly increased from April to May regardless of provenances (*p* <0.05, Figure 2, Supplemental material Appendix 7). The mass fraction of fibrous roots was slightly but significantly increased from April to May only in norther provenance (*p* <0.05, Figure 2). While there were not significant difference in average length of pioneer roots (Figure 3), it tended to increase from April to May in eastern provenance while decrease in northern provenance. On the other hand, the total number of pioneer roots was significantly increased only in northern provenance (*p* <0.05, Figure 3).

**Figure 2:**
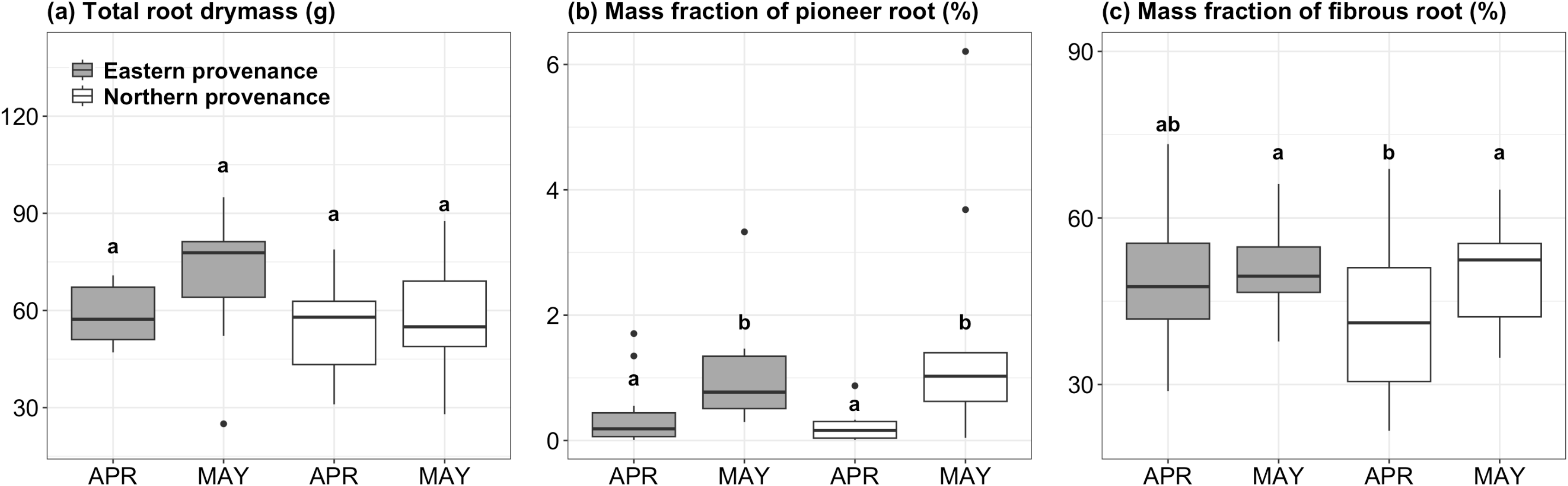
Mass variation at a whole root level between sampling months (APR: April, MAY: May) and provenances (grey: Eastern provenance, white: Northern provenance). Different letters denote significant difference in each panel (p <0.05).

**Figure 3:**
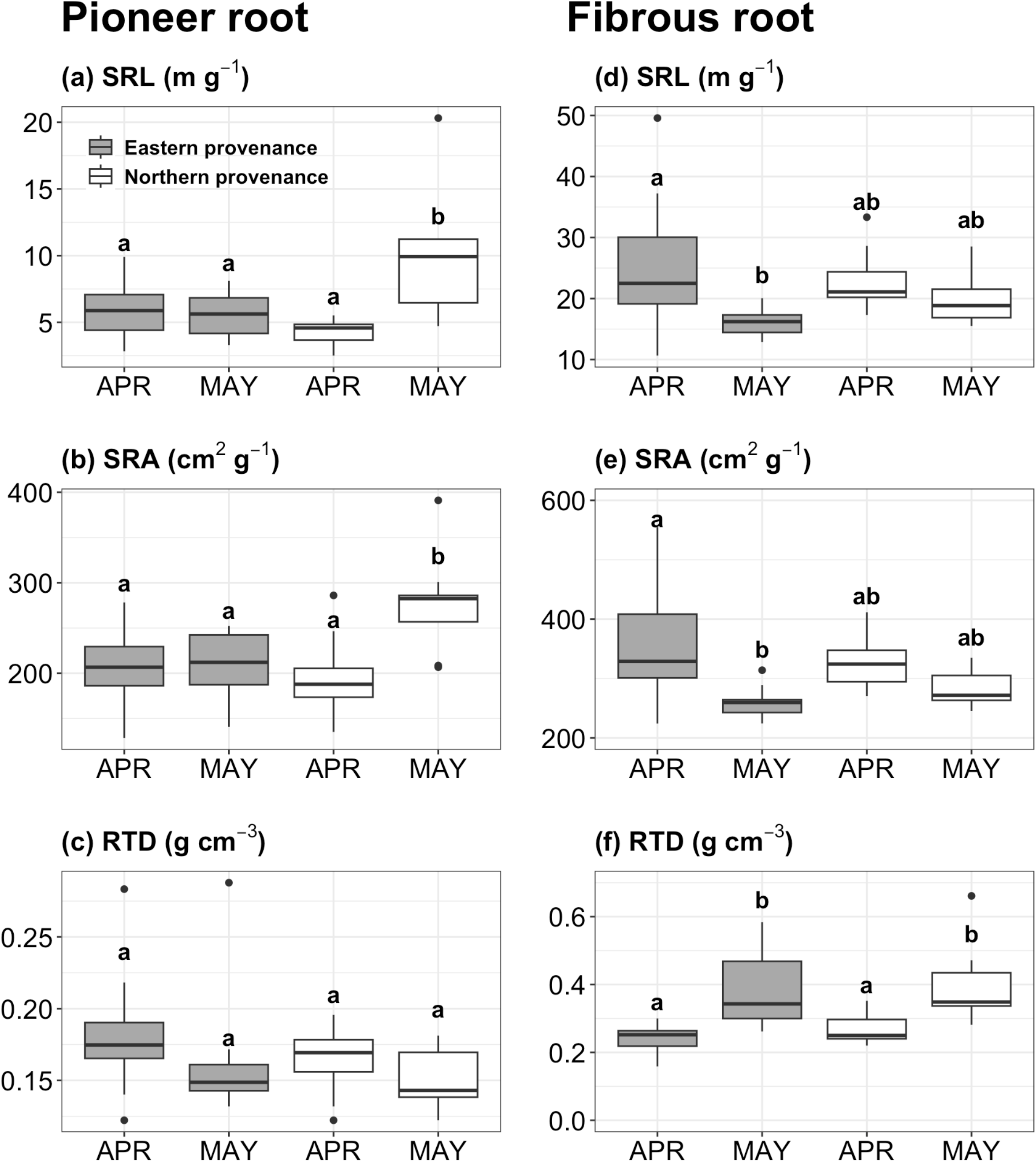
Variation in the average length (a) and total root number (b) of pioneer roots between sampling months and provenances.

In pioneer roots, significant increases of SRL and SRA from April to May were observed only in northern provenance (*p* <0.05, Figure 4, Supplemental material Appendix 8). Any significant changes of RTD were not observed in pioneer roots. In contrast, SRL and SRA of fibrous roots were significantly decreased from April to May only in eastern provenance (*p* <0.05, Figure 4). Contrary to pioneer roots, RTD of fibrous roots was significantly increased from April to May regardless of provenances (*p* <0.05, Figure 4, Supplemental material Appendix 8).

**Figure 4:**
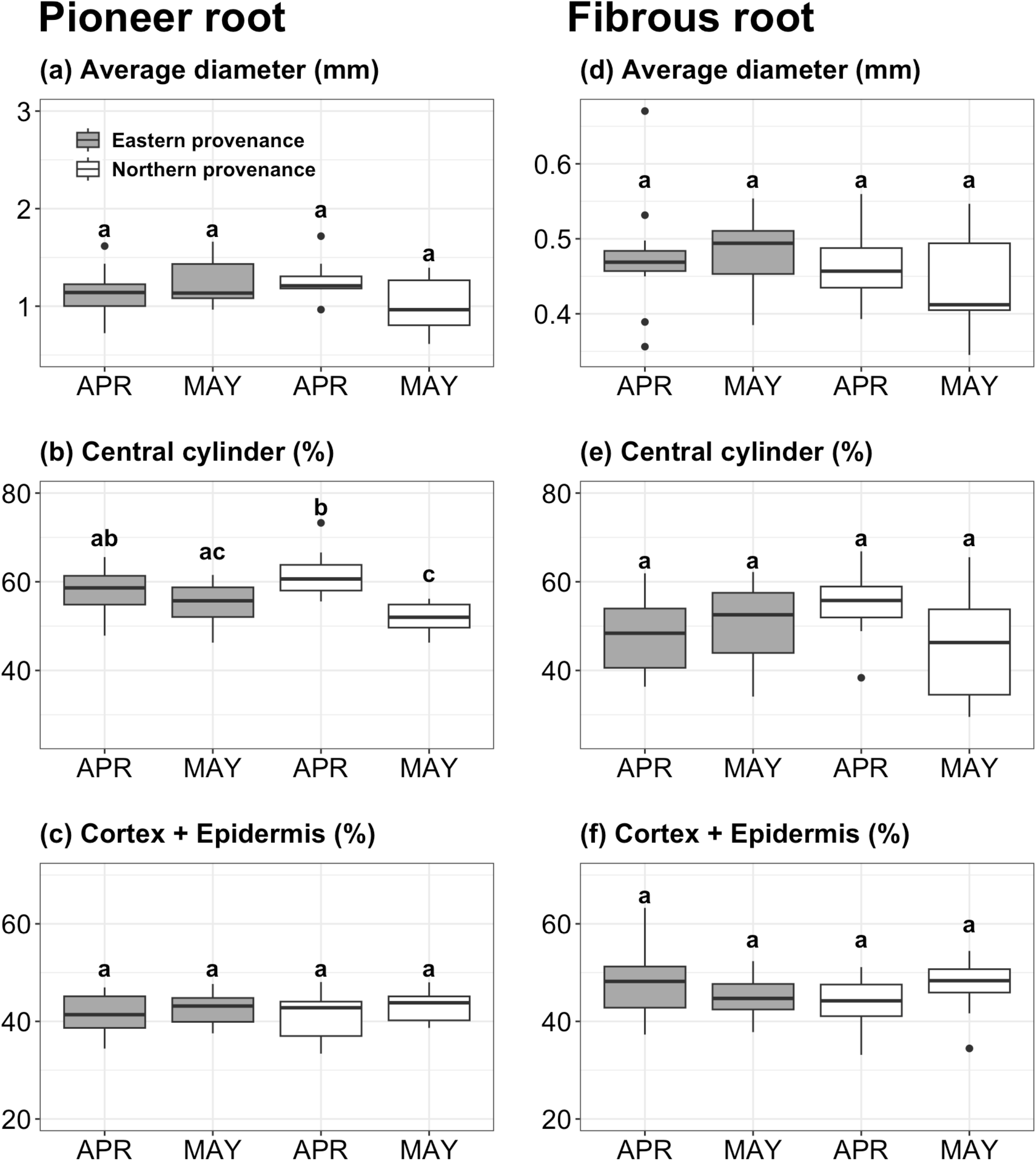
Morphological variation between sampling months and provenances in pioneer roots (a-c) and fibrous roots (d-f), respectively. SRL: specific root length, SRA: specific root area, RTD: root tissue density.

The interaction effect between sampling months and provenances was significant in average diameter of pioneer roots (*p* <0.05, Supplemental material Appendix 8). While there were not significant difference in average diameter of pioneer roots (Figure 5), its pattern of diameter change from April to May was similar to the average length (Figure 3), namely it tended to increase from April to May in eastern provenance while decrease in northern provenance. The interaction effect between sampling months and provenances was also significant in relative size of central cylinder of pioneer roots (*p* <0.05, Supplemental material Appendix 8). Only in northern provenance, the relative size of central cylinder was significantly decreased from April to May (*p* <0.05, Figure 5). In contrast, any significant change of average diameter, the relative size of central cylinder, and cortex and epidermis was not detected in fibrous roots.

**Figure 5:**
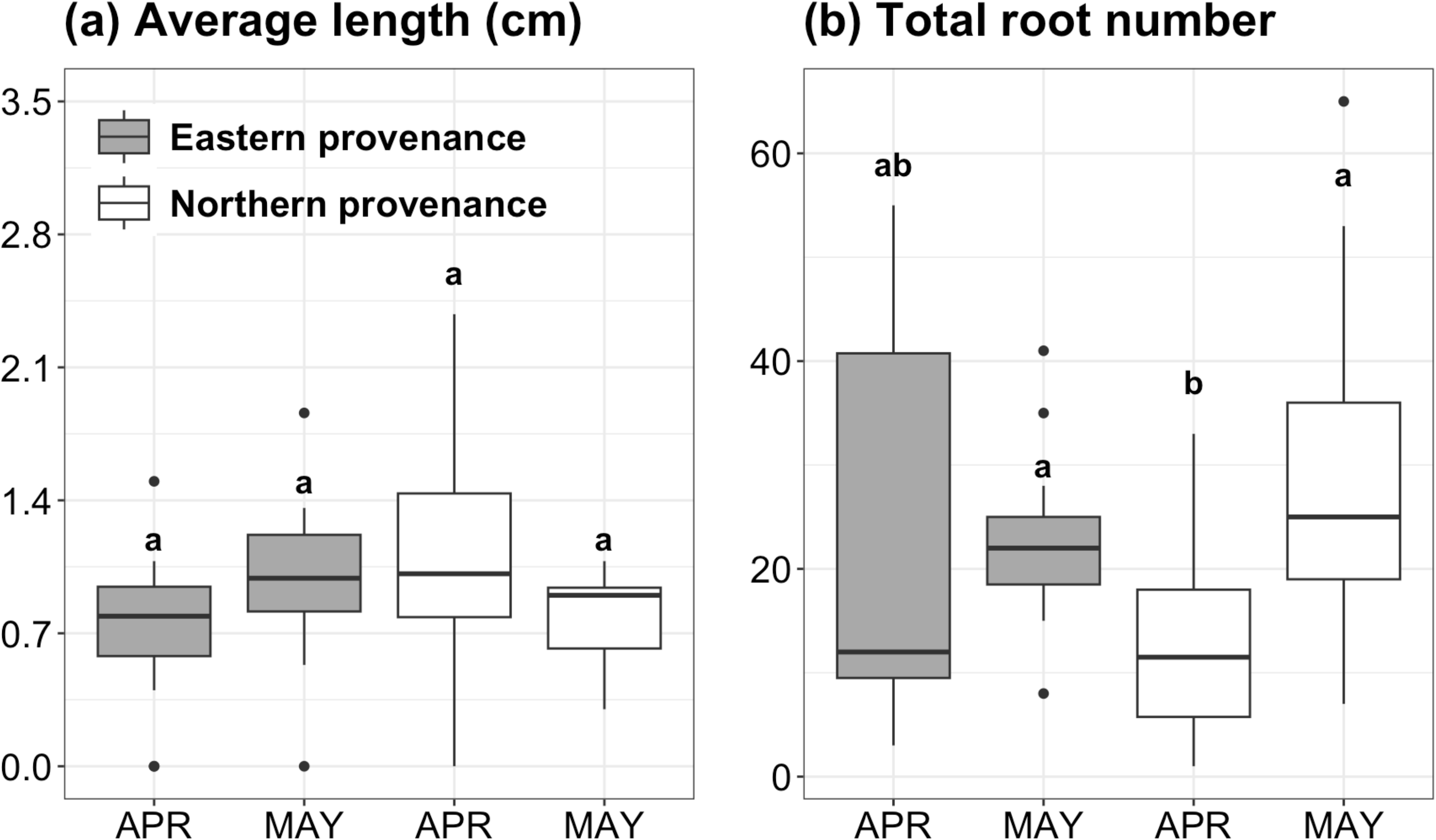
Average diameter and structural variation between sampling months and provenances in pioneer roots (a-c) and fibrous roots (d-f), respectively.

## Discussion

### Spring dynamics of pioneer roots

Previously, the quantitative dynamics of pioneer roots have been elucidated (Nakahata 2020, Montagnoli et al. 2021) with respect to the seasonal changes of the soil temperature (Ding et al. 2020), whereas relatively few studies have investigated the seasonal variation of construction costs and structures of pioneer roots (Zadworny & Eissenstat 2011). Only in the seedlings originating from heavy snow cover provenance (i.e., northern provenance), morphology and structure of pioneer roots significantly varied between April and May (Figure 4, Figure 5), while the quantity of pioneer roots was increased during the experimental periods regardless of provenances (Figure 2). To the best of our knowledge, this is the first empirical data regarding the intraspecific differences in spring growth dynamics of pioneer roots. These results would consist with previous studies (Satoh 1995) as well as our preliminary survey (Supplemental material Appendix 3). In fact, the increase in the number of root development in response to conditions of increasing soil temperature (Camm & Harper 1991). While root growth is fundamentally activated as soil temperature increases from winter to spring in areas with adequate precipitation (Pregitzer et al. 2000), it has been reported that the threshold of soil temperature at which roots can grow in conifers is approximately 0 to 4 °C (Wang et al. 2018, Ding et al. 2020). Beyond our expectation, however, the fact that the pioneer roots were observed even in April should raise the question as to when the pioneer roots of April emerged. At our experimental site, where the ground temperature remained near 0°C due to the thick snow cover and its insulation, the well-developed pioneer roots were observed (Supplemental material Appendix 3). It is thus possible that changes in the physical environment other than soil temperature, such as soil moisture conditions associated with snow thawing (Sanmiguel-Vallelado et al. 2021), and finer-scale nutrient conditions (Pregitzer et al. 2000) are relevant to spring root dynamics.

### Spring dynamics of fibrous roots

As soil nutrients are generally heterogeneous, choosing the location for nutrient absorption is important for plants to optimize the balance between construction costs of roots and benefits. Thus, the plasticity of fibrous roots is expected to have the ability to regulate construction costs (de Kroon & Visser 2003, Begon et al. 2006). In accordance with our hypothesis, RTD of fibrous roots in both provenances were increased from April to May (Figure 4). We also observed the non-significant changes in average diameter and other structural traits in May (Figure 5). These results can demonstrate the matured fibrous roots would develop from April to May. This would partly support the sensitivity to drought as well as soil temperature of fibrous roots (Ding et al. 2020). Although we did not directly evaluate soil water condition (Supplemental material Appendix 2), the soil was relatively dry in May, a month after the snow thawed out in April. It has been reported that high RTD would denote the stress resistance and long life of roots (Weemstra et al. 2016)

### Implications for the intraspecific variation in spring root dynamics

Understanding phenotypic development is synonymous with understanding its evolutionary processes (West-Eberhard 2003). The intraspecific variation due to local adaptation would represent a trade-off between growth and survival (Csilléry et al. 2020). In high snow cover regions, soil temperature and resource availability would increase rapidly (Bardgett et al. 2005) more than low snow cover regions because soil suddenly faced relatively high air temperatures when the thermal interception by snow cover ends (Fukuzawa et al. 2021, Makoto et al. 2022). As a result, after thawing of snow cover, the niche suddenly become available and competition for soil space and resources intensifies (Begon et al. 2006). In May, we observed the pioneer roots with lower construction costs and immature central cylinder, indicating that it would be the relatively young pioneer roots with inferior function of soil space exploration. This interception was further supported by the quantitative dynamics of pioneer roots (Figure 2, Figure 3), suggesting that the newly pioneer roots were developed from April with response to increasing relatively lower soil temperature, i.e., from below 8 °C, approximately (Wang et al. 2018). These results would support our hypothesis. Namely, the intraspecific variation in the ability to explore soil space may occurs between high snow cover regions with high competition and low snow cover regions with severe soil freezing.

On the other hand, in fibrous roots, SRL and SRA were significantly decreased from April to May only in eastern provenance (Figure 3), indicating the construction costs was low and the function of soil nutrients absorption was high in April. We suspect these results may be plastic responses in eastern provenance to the environment of common garden with high snow cover condition. This would be partly supported by the difference of root morphology between growing sites, i.e., the common garden and eastern provenance site (Supplemental material Appendix 6), where the average values (± standard error) of SRL and SRA in eastern provenance site were 18.9 ± 1.67 m g^-1^ and 320.79 ± 9.97cm^2^ g^-1^ in April. Although the statistical comparison was not able to be performed, the relatively higher values of these parameters were observed in common garden rather than a provenance site in April. This may be associated with the tolerance to soil freezing or low temperature of fibrous roots in eastern provenance (Sakai & Larcher 1987, Ambroise et al. 2020). Given that roots are the least freeze-tolerant of all plant organs and soil freezing can be a selection pressure (Ambroise et al. 2020), the contrasting snow cover environments may consequently be realized as the intraspecific variation in freezing tolerance in fibrous roots. If so, since the soil temperature in April at common garden may be relatively high for seedlings of eastern provenance, they may start the development of nutrient absorption even just after snow thawed out. In other aspect, recent studies have reported dramatic changes in soil microbial communities due to spring snow thawing (Broadbent et al. 2021), which was roughly common in the study site (Supplemental material Appendix 1). Although this may be the chance of nutrient absorption for eastern provenance, further experimental approach should be needed to verify whether there was the intraspecific variation in the tolerance to soil freezing or low temperature of fibrous roots.

## Conclusion

This study elucidated the spring growth dynamics of pioneer roots and fibrous roots in Sakhalin fir seedlings originating from provenances with contrasting snow cover environments. The obtained results provided the eco-evolutional insights for mechanistical processes in niche acquisition under variable snow cover environments in boreal regions. The difference in competitive versus conservative soil environments due to variable snow cover condition may be responsible for the intraspecific variation in root function to explore soil space. Contrary to our expectation, in low snow cover provenance, the intraspecific variation was observed in morphological traits in fibrous roots, perhaps reflecting the plastic responses to growing sites with high snow cover environments. Since we did not reveal the critical factors explaining why root developmental patterns varied between original provenances, future studies are expected to find the genetic evidence associated with the intraspecific variation of spring root dynamics. The insights should enhance the understanding of the adaptive phenology in trees to changing environments.

## Acknowledgements

We thank the staffs of Hokkaido Research Organization for maintaining the common garden trial and its plant materials and helping the measurement. The authors declare no conflicts of interest associated with this manuscript. The datasets analyzed during the current study are available from the corresponding author on reasonable request. TS conceived and managed the study project. WI provided and managed the experimental materials and filed. TS, WI, TM, IE, JI, MK, NM conducted the field samplings with the methodology of sampling and survey developed by MK and NM.

All authors analyzed the samples. TS statistically analyzed the data and wrote the draft of manuscript, and all other authors provided the editorial advice.

## References

Ambroise, V, Legay, S, Guerriero, G, Hausman, JF, Cuypers, A, Sergeant, K (2020) The roots of plant frost hardiness and tolerance. Plant and Cell Physiology, 61:3–20 doi: 10.1093/pcp/pcz196

Bardgett, RD, Bowman, WD, Kaufmann, R, Schmidt, SK (2005) A temporal approach to linking aboveground and belowground ecology. Trends in ecology & evolution, 20:634–641 doi: 10.1016/j.tree.2005.08.005

Begon, M, Townsend, CR, Harper, JL (2006) Resources. In Ecology: from individuals to ecosystems. Wiley, New York, Chapter 3, pp.65–101.

Broadbent, AA, Snell, HS, Michas, A, Pritchard, WJ, Newbold, L, Cordero, I, Goodall, T, Schallhart, N, Kaufmann, R, Griggiths, RI, Schloter, M, Bahn, M, Bardgett, RD (2021) Climate change alters temporal dynamics of alpine soil microbial functioning and biogeochemical cycling via earlier snowmelt. The ISME journal, 15:2264–2275 doi: 10.1038/s41396-021-00922-0

Camm, EL, Harper, GJ (1991) Temporal variations in cold sensitivity of root growth in cold-stored white spruce seedlings. Tree Physiology, 9:425–431 doi: 10.1093/treephys/9.3.425

Csilléry, K, Ovaskainen, O, Sperisen, C, Buchmann, N, Widmer, A, Gugerli, F (2020) Adaptation to local climate in multi-trait space: evidence from silver fir (*Abies alba* Mill.) populations across a heterogeneous environment. Heredity, 124:77–92 doi: 10.1038/s41437-019-0240-0

De Kort, H, Vander Mijnsbrugge, K, Vandepitte, K, Mergeay, J, Ovaskainen, O, Honnay, O (2016) Evolution, plasticity and evolving plasticity of phenology in the tree species *Alnus glutinosa*. Journal of evolutionary biology, 29:253–264 doi: 10.1111/jeb.12777

de Kroon, H, Visser, EJ (2003) Distribution of Roots in Soil, and Root Foraging Activity. In *Root ecology*. Springer, Berlin, Chapter 2, pp.33–60. doi: 10.1007/978-3-662-09784-7_2

De Lisle, SP, Mäenpää, MI, Svensson, EI (2022) Phenotypic plasticity is aligned with phenological adaptation on both micro and macroevolutionary timescales. Ecology letters, 25:790–801 doi: 10.1111/ele.13953

Ding, Y, Schiestl-Aalto, P, Helmisaari, HS, Makita, N, Ryhti, K, Kulmala, L (2020) Temperature and moisture dependence of daily growth of Scots pine (*Pinus sylvestris* L.) roots in Southern Finland. Tree Physiology, 40:272–283 doi: 10.1093/treephys/tpz131

Eiga, S (1984) Ecological study on the freezing resistance of Saghalin fir (*Abies sachalinensis* Mast.) in Hokkaido. Bull. For. Tree Breed. Inst. 2:61–107 (in Japanese with English abstract)

Ellington, EH, Bastille Rousseau, G, Austin, C, Landolt, KN, Pond, BA, Rees, EE, Robar, N, Murray, DL (2015) Using multiple imputation to estimate missing data in meta regression. Methods in Ecology and Evolution, 6:153–163. doi: 10.1111/2041-210X.12322

Forrest, J, Miller-Rushing, AJ (2010) Toward a synthetic understanding of the role of phenology in ecology and evolution. Philosophical Transactions of the Royal Society B: Biological Sciences, 365:3101–3112 doi: 10.1098/rstb.2010.0145

Franks, SJ, Weber, JJ, Aitken, SN (2014) Evolutionary and plastic responses to climate change in terrestrial plant populations. Evolutionary applications, 7:123–139 doi: 10.1111/eva.12112

Fréchette E, Chang CYY, Ensminger I (2020) Variation in the phenology of photosynthesis among eastern white pine provenances in response to warming. Global Change Biology, 26:5217–5234 doi: 10.1111/gcb.15150

Fukuzawa, K, Tateno, R, Ugawa, S, Watanabe, T, Hosokawa, N, Imada, S, Shibata, H (2021) Timing of forest fine root production advances with reduced snow cover in northern Japan: implications for climate-induced change in understory and overstory competition. Oecologia, 196:263–273 doi: 10.1007/s00442-021-04914-x

Ghalambor, CK, McKay, JK, Carroll, SP, Reznick, DN (2007) Adaptive versus non adaptive phenotypic plasticity and the potential for contemporary adaptation in new environments. Functional ecology, 21:394–407 doi: 10.1111/j.1365-2435.2007.01283.x

Ishizuka, W, Kon, H, Kita, K, Kuromaru, M, Goto, S (2021) Local adaptation to contrasting climatic conditions in Sakhalin fir (*Abies sachalinensis*) revealed by long-term provenance trials. Ecological Research, 36:720–732 doi: 10.1111/1440-1703.12232

Liu, H, Wang, H, Li, N, Shao, J, Zhou, X, van Groenigen, KJ, Thakur, MP (2022) Phenological mismatches between above-and belowground plant responses to climate warming. Nature climate change, 12:97–102 doi: 10.1038/s41558-021-01244-x

Ma, T, Parker, T, Unger, S, Gewirtzman, J, Fetcher, N, Moody, ML, Tang, J (2022) Responses of root phenology in ecotypes of *Eriophorum vaginatum* to transplantation and warming in the Arctic. Science of The Total Environment, 805, 149926 doi: 10.1016/j.scitotenv.2021.149926

Makoto, K, Wilson, SD, Sato, T, Blume Werry, G, Cornelissen, JH (2020) Synchronous and asynchronous root and shoot phenology in temperate woody seedlings. Oikos, 129:643–650 doi: 10.1111/oik.06996

Makoto, K, Templer, PH, Katayama, A, Seki, O, Takagi, K (2022) Early snowmelt by an extreme warming event affects understory more than overstory trees in Japanese temperate forests. Ecosphere, 13:e4182 doi: 10.1002/ecs2.4182

McCormack, ML, Gaines, KP, Pastore, M, Eissenstat, DM (2015) Early season root production in relation to leaf production among six diverse temperate tree species. Plant and Soil, 389:121–129 doi: 10.1007/s11104-014-2347-7

Mediavilla, S, Escudero, A (2009) Ontogenetic changes in leaf phenology of two co-occurring Mediterranean oaks differing in leaf life span. Ecological Research, 24:1083–1090 doi: 10.1007/s11284-009-0587-4

Nakahata, R (2020) Pioneer root invasion and fibrous root development into disturbed soil space observed with a flatbed scanner method. Trees, 34:731–743 doi: 10.1007/s00468-020-01953-4

Nissen, J, Donatello, R, Van Dusen, B (2019) Missing data and bias in physics education research: A case for using multiple imputation. Physical Review Physics Education Research, 15:020106. doi: 10.1103/PhysRevPhysEducRes.15.020106

Noordermeer, D, Velasco, V, Ensminger, I (2021) Autumn warming delays downregulation of photosynthesis and does not increase risk of freezing damage in interior and coastal Douglas-fir seedlings. Frontiers in Forest Global Change, 4:688534 doi: 10.3389/ffgc.2021.688534

Persson, H (1978) Root dynamics in a young Scots pine stand in Central Sweden. Oikos, 30:508–519 doi: 10.2307/3543346

Pregitzer, KS, King, JS, Burton, AJ, Brown, SE (2000) Responses of tree fine roots to temperature. New Phytologist, 147:105–115 doi: 10.1046/j.1469-8137.2000.00689.x

R Core Team (2019) R development core team. R: a language and environment for statistical computing, 55:275–286.

Sakai, A (1983) Comparative study on freezing resistance of conifers with special reference to cold adaptation and its evolutive aspects. Canadian Journal of Botany, 61:2323–2332 doi: 10.1139/b83-255

Sakai, A, Larcher, W (1987) Frost Resistance in Plants. In: Frost Survival of Plants. Springer, Berlin, Chapter 6, pp.138–173.

Sanmiguel-Vallelado, A, Camarero, JJ, Morán-Tejeda, E, Gazol, A, Colangelo, M, Alonso-González, E, López-Moreno, JI (2021) Snow dynamics influence tree growth by controlling soil temperature in mountain pine forests. Agricultural and Forest Meteorology, 296:108205 doi: 10.1016/j.agrformet.2020.108205

Satoh, T (1995) Fundamental studies on the root growth of trees in Hokkaido. Bulletin of the Hokkaido Forestry Research Institute, 31:1–54. (in Japanese with English abstract)

Steele, SJ, Gower, ST, Vogel, JG, Norman, JM (1997) Root mass, net primary production and turnover in aspen, jack pine and black spruce forests in Saskatchewan and Manitoba, Canada. Tree physiology, 17:577–587 doi: 10.1093/treephys/17.8-9.577

Sugai, T, Ishizuka, W, Watanabe, T (2023) Landscape gradient of autumn photosynthetic decline in *Abies sachalinensis* seedlings. Journal of Forestry Research, 34:187–195 doi: 10.1007/s11676-022-01592-0

Tamura, A, Oguma, H, Fujimoto, R, Kuribayashi, M, Makita, N (2022) Phenology of fine root and shoot using high frequency temporal resolution images in a temperate larch forest. Rhizosphere, 22, 100541 doi: 10.1016/j.rhisph.2022.100541

Wang, Y, Mao, Z, Bakker, MR, Kim, JH, Brancheriau, L, Buatois, B, Leclerc, R, Selli, L, Rey, H, Jourdan, C, Stokes, A (2018) Linking conifer root growth and production to soil temperature and carbon supply in temperate forests. Plant and Soil, 426:33–50 doi: 10.1007/s11104-018-3596-7

Weemstra, M., Mommer, L., Visser, EJ, van Ruijven, J., Kuyper, TW, Mohren, GM, Sterck, FJ (2016) Towards a multidimensional root trait framework: a tree root review. New Phytologist, 211:1159–1169. doi: 10.1111/nph.14003

West-Eberhard, MJ (2003) Development. In: Developmental plasticity and evolution. Oxford University Press, Oxford, Chapter 5, pp.89–90.

Wilcox, H (1954) Primary organization of active and dormant roots of noble fir, *Abies procera*. American Journal of Botany, 41:812–821 doi: 10.2307/2438547

Yahara, H, Tanikawa, N, Okamoto, M, Makita, N (2019) Characterizing fine-root traits by species phylogeny and microbial symbiosis in 11 co-existing woody species. Oecologia, 191, 983–993 doi: 10.1007/s00442-019-04546-2

Zadworny, M, Eissenstat, DM (2011) Contrasting the morphology, anatomy and fungal colonization of new pioneer and fibrous roots. New Phytologist, 190:213–221 doi: 10.1111/j.1469-8137.2010.03598.x

