## Supplemental material Appendix for "Spring growth variation in pioneer and fibrous roots in *Abies sachalinensis* seedlings from provenances with contrasting snow cover environments"

### Appendix 1. Provenance environmental conditions

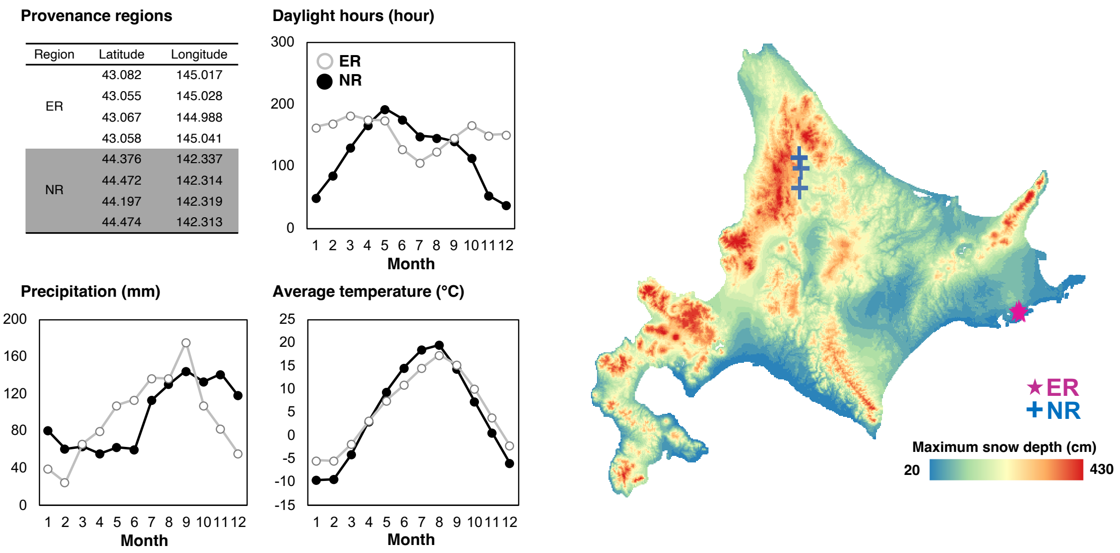


(Left) The latitude and longitude of mother trees for eastern (ER) and northern regions (NR), and its seasonal changes and of 40-year average of several meteorological values from 1981 to 2020. (Right) 1×1 km mesh map for the 40-year average of maximum snow depth, where the locations of mother trees for ER (red stars) and NR (blue cross) were indicated. Meteorological data was obtained from the Agro-Meteorological Grid Square Data, NARO (https://amu.rd.naro.go.jp/).

**Appendix 2. Growing environmental conditions in the experimental site**

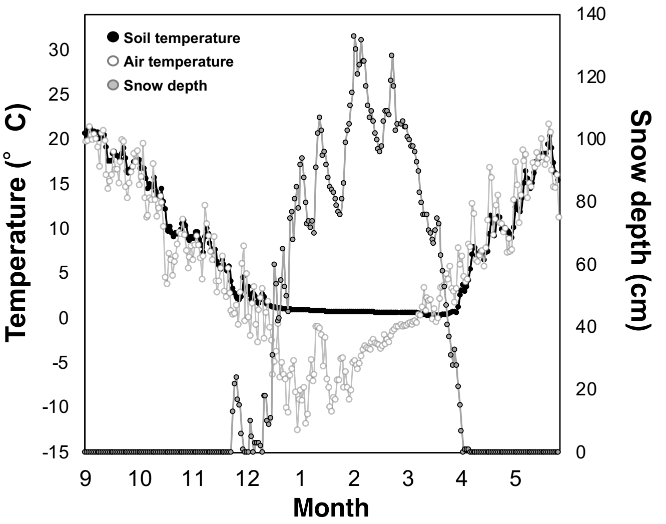


| Soil properties | Depth | April | May |
|---|---|---|---|
| pH (KCl) | 0 - 5 cm | 4.36 (0.06) | 4.54 (0.15) |
| | 5 - 15 cm | 4.35 (0.21) | 4.44 (0.17) |
| | 15 - 30 cm | 4.47 (0.17) | 4.49 (0.14) |
| EC (KCl, mS cm$^{-1}$) | 0 - 5 cm | 79.85 (0.88) | 66.33 (1.27) |
| | 5 - 15 cm | 88.52 (0.88) | 72.02 (1.56) |
| | 15 - 30 cm | 84.31 (0.82) | 70.13 (0.54) |

(Left) Seasonal changes of daily averaged temperature of soil (black) and air (white), and snow depth (grey) from September 1, 2021, to May 31, 2022 at the experimental site. Soil temperature was monitored in the ground at a depth of 10–20 cm, using a data logger (TW-71NW, T&D Corp., Japan) with a thermos recorder steel probe (TR-0406, T&D Corp., Japan). Air temperature was monitored at ca. 30 cm above the ground not in a winter period, and ca. 1 m in winter periods from December to April, by a data logger (HOBO UA-002-64, Onset Computer Corp., Bourne, MA, USA). Snow depth data was obtained from the Agro-Meteorological Grid Square Data, NARO (https://amu.rd.naro.go.jp/). (Right) Soil chemical properties in each sampling month at the experimental site. Soil pH was measured by a portable pH meter (pH-33B, HORIBA Advanced Techno, Co., Ltd., Tokyo, Japan). Soil EC was measure by a portable EC meter (EC-33B, HORIBA Advanced Techno, Co., Ltd., Tokyo, Japan). The soil was air-dried and sieved through a 2 mm mesh. The solution measured was adjusted at specific ratios (1: 5 = dry soil mass: 1M KCl). 1 ml of the supernatant suspension was used for the measurements. Measurements were taken twice for each sample and the average value was used.

**Appendix 3. Winter root sampling and its appearances**

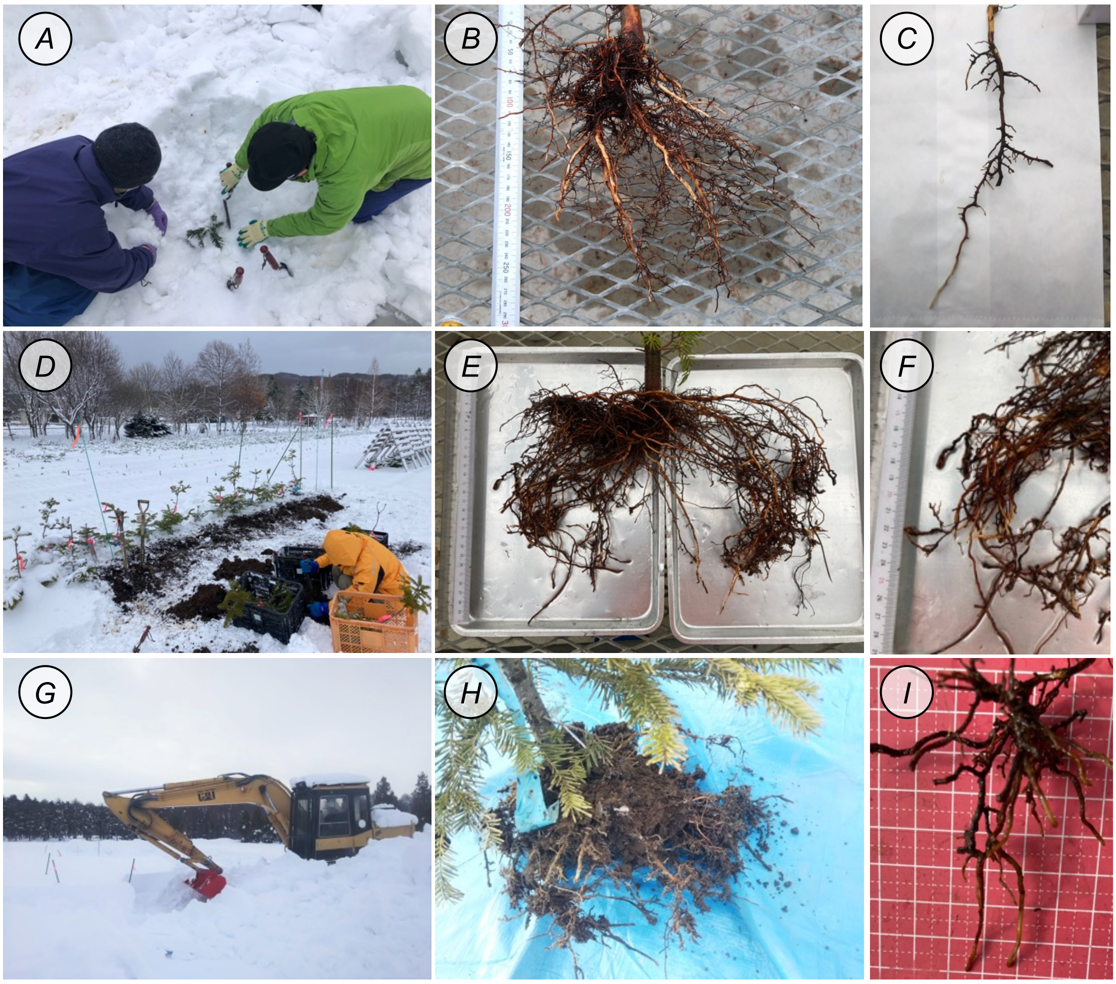


Root from the seedlings covered by the snow were collected in March (*A* - *C*) as a preliminary experiment, December 2021 (*D* - *F*), and February 2022 (*G* - *I*) as post experiments. In the all sampling months, the pioneer roots were observed in *Abies sachalinensis* seedlings.

**Appendix 4. Appearance of pioneer roots in *Abies Sachalinensis***

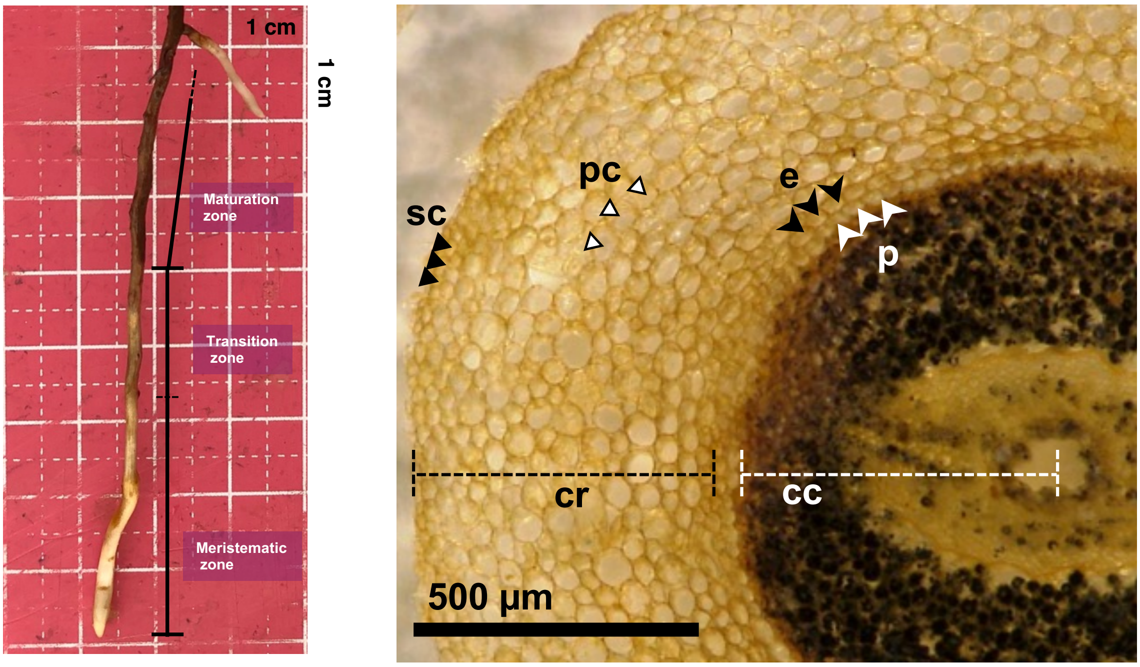


(Left) Longitudinal tissue structures recognizable from the color of pioneer roots and (Right) its anatomical phenotypes in *Abies Sachalinensis* seedlings. Abbreviations for anatomical phenotypes are following; e: endodermis, p: pericycle, cc: central cylinder, cr: cortex, pc: parenchymal cell, sc: sclerenchyma cell.

**Appendix 5. Difference between fibrous and lignified woody roots in *Abies sachalinensis***

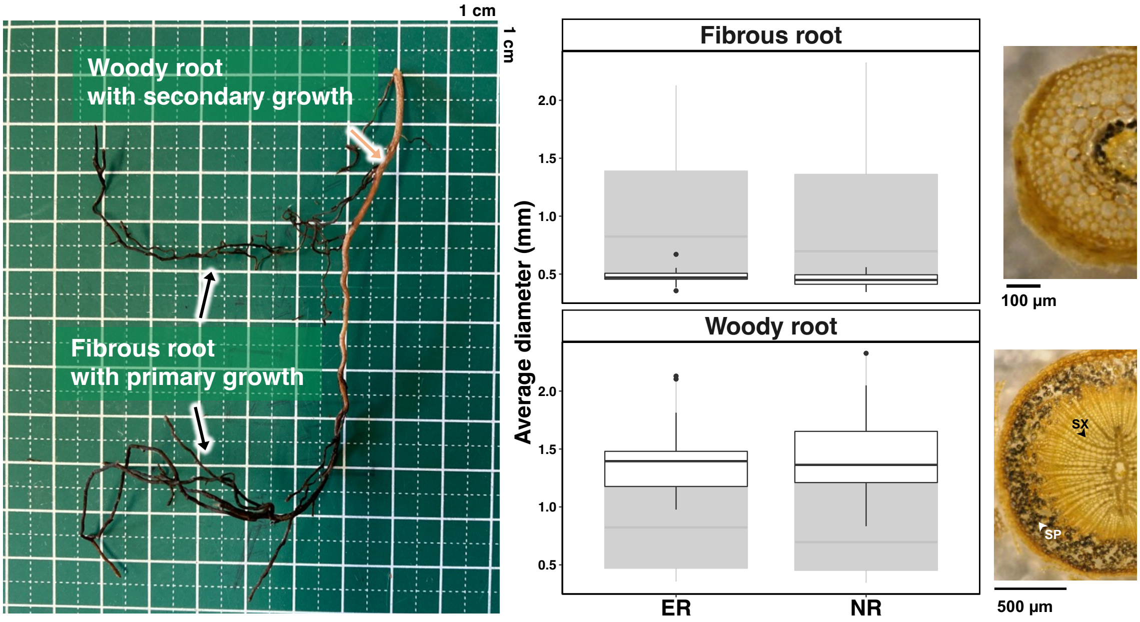


(Left) The appearance of fibrous and woody roots, (Center) their difference of average diameter between provenance eastern (ER) and northern regions (NR) with highlighted by root types, and (Right) its anatomical appearance in *Abies Sachalinensis* seedlings. While fibrous roots exhibit a brown dark color with relatively thin and especially soft, woody roots presents a light brown color with thick. Based on the section images, the roots collected as woody roots demonstrated secondary growth structures, such as secondary xylem (sx) and secondary phloem tissues (sp).

**Appendix 6. Morphological variation of fibrous roots in eastern provenance**

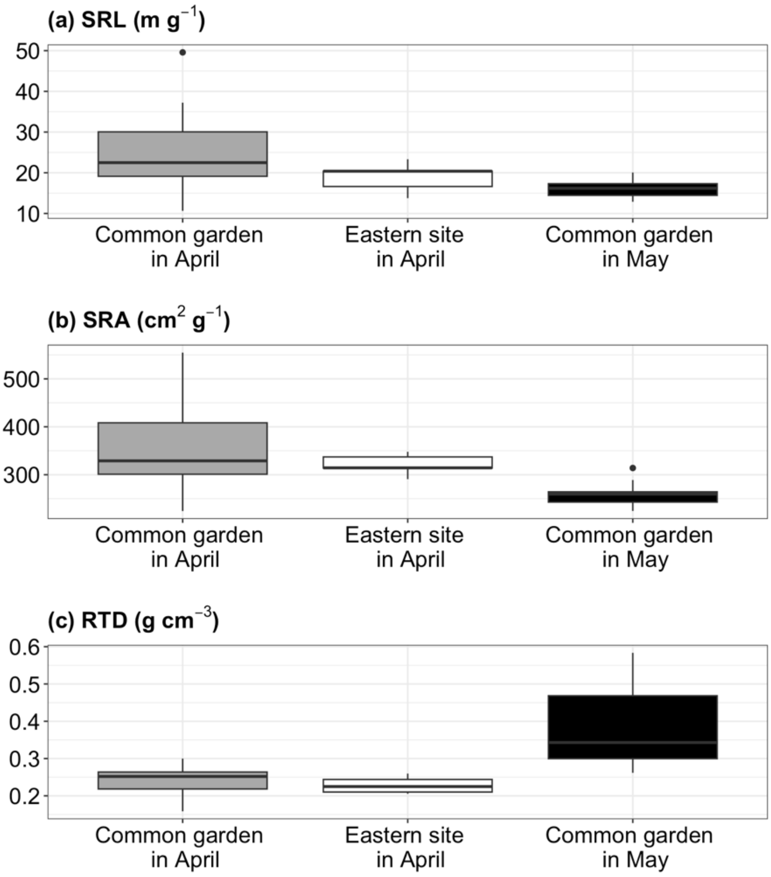


Morphological variation between sampling months and growing sites in fibrous roots of seedlings originating from eastern provenance, respectively. Five seedings were collected in an eastern provenance site (43°02′N, 145°02′E) the day before the collection date in the common garden (n=5). SRL: specific root length, SRA: specific root area, RTD: root tissue density.

**Appendix 7**. SUMMARY OF ANOVA FOR WHOLE ROOT AND PIONEER ROOTS

(*F*-value | *P*-value)

| Phenotype | Month (M) | Genetic (G) | M × G |
|---|---|---|---|
| Total root dry mass (g) | 1.80 \| 0.189 | 0.51 \| 0.486 | 0.52 \| 0.475 |
| Mass fraction of pioneer root (%) | 23.31 \| **<0.001** | 0.22 \| 0.647 | 0.18 \| 0.671 |
| Mass fraction of fibrous root (%) | 5.53 \| **<0.05** | 3.31 \| 0.096 | 0.77 \| 0.385 |
| Average length of pioneer roots (cm) | 2.58 \| 0.117 | 4.53 \| **<0.05** | 4.53 \| **<0.05** |
| Total number of pioneer roots | 5.71 \| **<0.05** | 2.08 \| 0.169 | 1.73 \| 0.197 |

NOTE.—Number denote F value in left and P value in right. Bold denote significant effects of sampling month (month, M), provenances (genetic, G), and its interaction.

**Appendix 8**. SUMMARY OF ANOVA AT INDIVIDUAL ROOTS LEVELS

(*F*-value | *P*-value)

| | | Pioneer roots | | | Fibrous roots | | |
|---|---|---|---|---|---|---|---|
| Level | Phenotype | Month (M) | Genetic (G) | M × G | M | G | M × G |
| Morphology | SRL (m $g^{-1}$) | 0.00 \| 0.948 | 2.89 \| 0.106 | **11.9 \| <0.01** | **15.6 \| <0.001** | 0.34 \| 0.564 | 2.92 \| 0.097 |
| | SRA ($cm^2$ $g^{-1}$) | 0.10 \| 0.749 | 0.29 \| 0.593 | **7.85 \| <0.01** | **21.0 \| <0.001** | 1.21 \| 0.29 | 2.97 \| 0.094 |
| | RTD (g $cm^3$) | 2.48 \| 0.124 | 1.42 \| 0.248 | 0.11 \| 0.736 | **20.3 \| <0.001** | 1.48 \| 0.238 | 0.25 \| 0.620 |
| | Average diameter (mm) | 1.06 \| 0.309 | 1.78 \| 0.197 | **7.25 \| <0.05** | 0.06 \| 0.807 | 0.09 \| 0.763 | 0.85 \| 0.363 |
| Anatomy | Central cylinder (%) | 3.99 \| 0.053 | 2.75 \| 0.114 | **6.09 \| <0.05** | 0.12 \| 0.727 | 2.84 \| 0.109 | 3.95 \| 0.055 |
| | Cortex + Epidermis (%) | 0.64 \| 0.426 | 0.00 \| 0.993 | 0.02 \| 0.882 | 1.00 \| 0.322 | 1.78 \| 0.202 | 3.16 \| 0.084 |

NOTE.—Number denote F value in left and P value in right. Bold denote significant effects of sampling month (month, M), provenances (genetic, G), and its interaction. SRL: specific root length, SRA: specific root area, RTD: root tissue density.
